# A positional information gradient of sonic hedgehog is required for flight feather formation in avian wings

**DOI:** 10.1101/698555

**Authors:** Lara Busby, Cristina Aceituno, Caitlin McQueen, Constance A. Rich, Maria A. Ros, Matthew Towers

**Affiliations:** Department of Biomedical Science, University of Sheffield, Western Bank, Sheffield, S10 2TN, UK; Instituto de Biomedicina y Biotecnología de Cantabria, IBBTEC (CSIC-Universidad de Cantabria). 39011 Santander, Spain; Departamento de Anatomía y Biología Celular. Facultad de Medicina. Universidad de Cantabria. 39011 Santander, Spain; Department of Genetics, University of Cambridge, UK; University of Florida, Gainesville, USA

## Abstract

Flight is a triumph of evolution that enabled the radiation and success of birds. A crucial step was the development of forelimb flight feathers that may have evolved for courtship or territorial displays in ancestral theropod dinosaurs. Classical tissue recombination experiments performed in the chick embryo provide evidence that signals operating during early limb development specify the position and identity of feathers. Here we show that a positional information gradient of Sonic hedgehog (Shh) signalling in the embryonic chick wing bud specifies the pattern of adult flight feathers in a defined spatial and temporal sequence that reflects their different identities. We reveal that the Shh signalling gradient is interpreted into specific patterns of flight feather-associated gene expression. Our data suggests that flight feather evolution involved the co-option of the pre-existing digit patterning mechanism and therefore uncovers an embryonic process that played a fundamental step in the evolution of avian flight.

## Introduction

Flight feathers provide most of the flapping, gliding and soaring ability required for airborne locomotion in birds. Three identities of flight feather—based on differences in size, shape and location—are present in bird wings: primaries along the posterior margin of digits 2 and 3, secondaries along the posterior margin of the ulna, and alulars along the posterior margin of digit 1 ^1^ (Figs. 1a and b). Flight feathers are much longer and more rigid than other feathers, including covert feathers that adorn most of the surface of a bird, and down feathers that lie close to the body to provide insulation ^1^. They are also unique, both in being bilaterally asymmetric across the central midvein, and in forming strong ligamentous connections with the skeleton, which aids their independent movement during flight ^1,2^ (Fig. 1b—primaries to digits 2 and 3; secondaries to the ulna; alulars to digit 1). The presence of ‘flight feathers’ on the posterior margins of forearms of flightless bipedal theropod dinosaurs provides strong evidence that they evolved to fulfil another function such as defence or courtship, thus making it likely that they played an early and important step in the evolution of flight in later birds ^3–5^.

**Figure 1.**
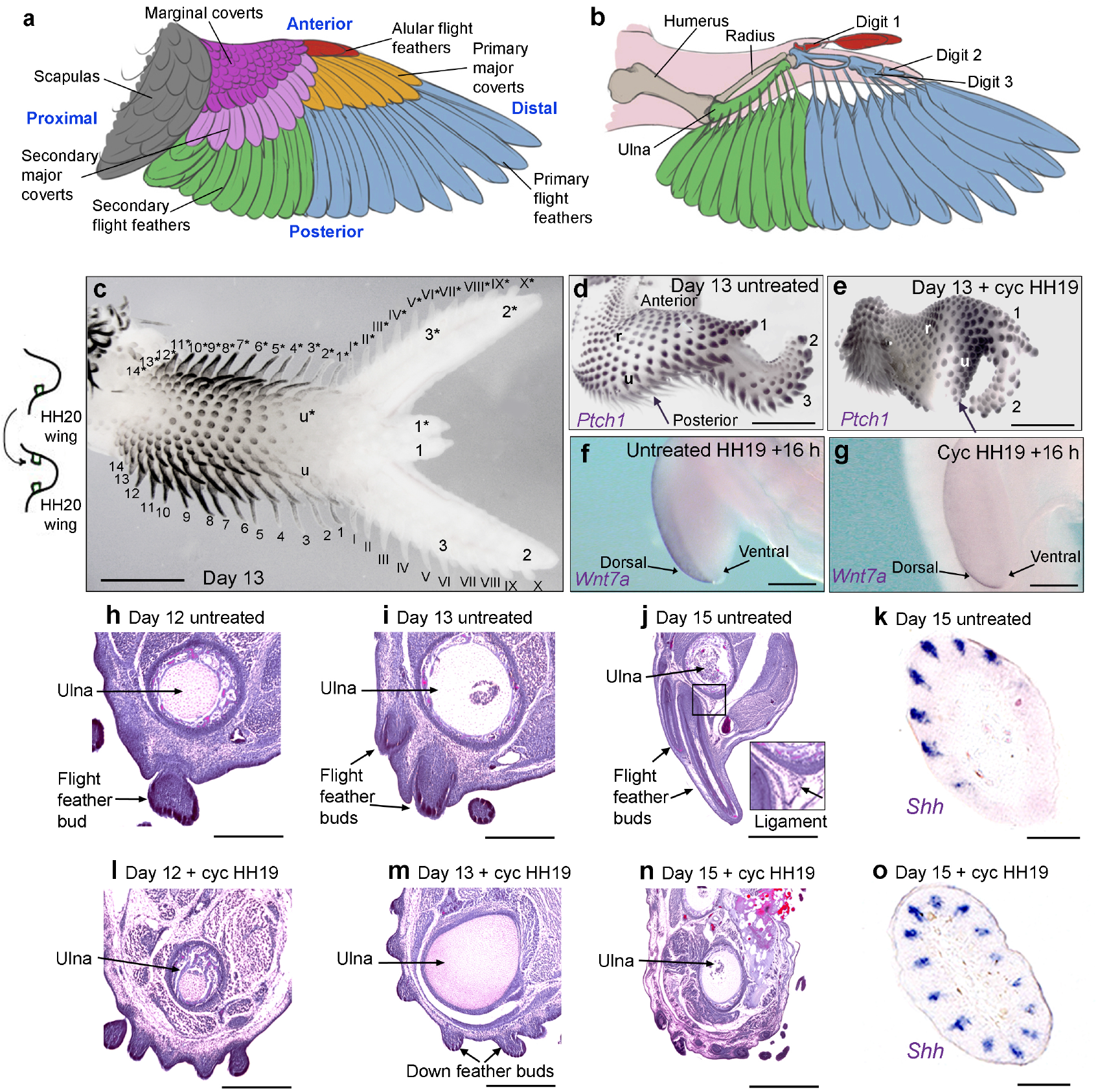
Shh signalling is required for flight feather bud formation. Schematics showing general bird wing feather pattern (**a**) and relationship with the skeleton (**b**). Graft of a HH20 polarising region made to the anterior of a host wing bud duplicates the flight feather pattern (**c**—Secondaries—Roman numerals, Primaries—Arabic numerals, note feather pigmentation). Developing flight feathers as shown by *Ptch1* in all buds in untreated day 13 wings (arrow—**d**), but not in wings of embryos treated with cyclopamine (cyc) at HH19 (arrow—*n*=4/4—**e**). Note—treatments at HH19 often result in loss of digit 3, but not the radius (r) or ulna (u) ^14^. *Wnt7a* expression in the dorsal epithelium of untreated (*n*=4/4—**f**) and HH19 cyclopamine-treated wing buds after 16 h (*n*=4/4—**g**). H&E staining on transverse sections of day 12 to day 15 forewings showing developing flight feathers in untreated (*n*=15/15—**h-j**) but not cyclopamine-treated wings (*n*=18/23—**l-n**). Asymmetric expression of *Shh* in flight feather buds in forewings of untreated wings (*n*=6/6 wings—**k**) but not in cyclopamine-treated wings ((*n*=6/6 wings—**o**). Scale bars: c-e 2 mm; f, g, j, n—150 μm; h, l—100 μm; i, m—125 μm; k, o—50 μm

Although much is known about the molecular pathways involved in the induction, positioning and morphogenesis of feathers ^6^, little is known about how different types of feathers are specified. The analyses of naturally occurring mutants provides evidence of a genetic pathway for flight feather development ^7,8^. In support of this proposal, it was recently revealed that the gene encoding the Sim1 transcription factor is a specific marker of the flight feather-forming regions of the bird wing ^9^. In addition, classical tissue recombination experiments in chickens provide evidence that signals acting at the earliest stages of wing bud development at around day 3 of incubation (HH20—see methods for staging) specify feather position and identity ^10,11^. An important signal known to function at this stage is Sonic hedgehog (Shh)—a protein that emanates from a transient signalling centre called the polarising region (also known as the zone of polarising region activity), which is located at the posterior margin of the limb bud ^12^. Shh is involved in the specification of antero-posterior positional values in the chick limb in a concentration-dependent manner between HH19 and HH22 ^13 14^ (thumb to little finger) and in stimulating proliferative growth along this axis ^15^ (reviewed in ^16^). However, it is unclear if Shh has a direct role in the specification of feather pattern, because although grafts of polarising region cells made to the anterior margin of host HH20 wing buds at incubation day 3.5 can duplicate all feather buds as shown at day 13 (Fig. 1c—primaries flight feathers in Roman numerals, secondaries in Arabic numerals), this could be an indirect consequence of all tissues being duplicated across the antero-posterior axis (Fig. 1c—ulna, digits 1, 2 and 3) ^17^.

In this study, we show that Shh signalling by the embryonic chick wing polarising region is required for the specification and formation of adult flight feathers. Our data provide evidence that a positional information gradient of Shh signalling integrates digit and flight feather patterning, and thus provides insights into the co-evolution of these important structures.

## Results

### Shh signalling is required for flight feather bud formation

During our extensive analyses of chick embryos in which the Shh signalling pathway was transiently inhibited with cyclopamine at day 3 of incubation (HH19/20) for approximately 72 h ^18^, we often noticed abnormal flight feather bud development. Thus, raised flight feather buds expressing *Ptch1*—a direct target of Shh signaling that is involved in feather morphogenesis ^19–21^—are found along the posterior margin of untreated day 13 wings (arrow in Fig. 1d), but not in the wings of embryos which were systemically treated with cyclopamine at HH19 (arrow in Fig. 1e), in which all of the feather buds have a similar morphology. Therefore, this observation demonstrates that it is the earlier loss of Shh signalling by the polarising region that perturbs flight feather bud formation, rather than the loss of Shh signalling within the buds themselves.

Flight feather buds form along the dorsal-ventral boundary of the wing, which if disrupted can result in abnormal flight feather bud development ^22^. Therefore, to examine if the loss of Shh signalling affects dorso-ventral patterning of the wing bud, we examined the expression of *Wnt7a*, which is expressed in the dorsal epithelium. In both the wing buds of untreated (arrows in Fig. 1f) and HH19 cyclopamine-treated embryos, the expression of *Wnt7a* reveals that the dorsal-ventral boundary remains intact after 16 h (arrows in Fig. 1g). Therefore, abnormal flight feather bud formation following the earlier transient loss of Shh signalling is not a consequence of defective dorso-ventral patterning.

Developing flight feather buds become morphologically distinct during late incubation stages by growing inwards to make ligamentous connections with the skeleton, and by displaying bilateral asymmetry ^1 2^. Therefore, we used these morphological characteristics to explore if Shh signalling is required for advanced stages of flight feather bud development. H&E staining on transverse sections of untreated forewings reveal that flight feather buds grow away from the posterior margin of the wing, and also invaginate into deeper tissues until they reach the ulna by day 13 ^2^ (Figs. 1h-j). In addition, developing flight feather buds can also be identified in transverse section at day 15 by their asymmetric pattern of *Shh* expression ^2^ (Fig. 1k). However, in embryos treated with cyclopamine at HH19, flight feather buds frequently fail to form along the posterior border of the wing, although it remains covered with natal down buds, none of which invaginate deeply towards the skeleton (Figs. 1l-n—Supplementary Table 1). Furthermore, only developing feather buds with symmetric expression of *Shh* are observed in forewing regions of wings at day 15, thus again demonstrating that flight feather buds are selectively missing (Fig. 1o).

Therefore, these observations demonstrate a transient and specific requirement for Shh signalling by the polarising region for later flight feather bud development.

### Shh signalling is required for flight feather-associated gene expression

Recently, molecular markers of the flight-feather forming regions of the chick wing have been identified which include *Sim1* ^9^ and *Zic1* ^23^. Notably, *Sim1* has an avian-specific forelimb expression pattern in the dermis ^24 9^. Therefore, to examine if the inhibition of Shh signalling affects flight feather bud-associated gene expression, we performed a series of RNA sequencing experiments on tissue dissected from day 10 wings. This stage was selected because it is when flight feather buds become morphologically distinct from other feather buds (see expression of a general marker, *Bmp7* ^25^, in raised flight feather buds at this stage—Supplementary Figure 1). We also sequenced RNA from soft tissue flanking the posterior margin of the ulna that forms normally in all wings treated with cyclopamine at HH19.

We contrasted sequencing data from the posterior forewing regions of cyclopamine-treated and untreated Bovans brown wings (Fig. 2a; top ten up- and down-regulated genes by >5-fold shown—Supplementary Information). We also enriched for genes associated with feather bud development by contrasting RNA-seq datasets obtained from Bovans brown wings with datasets obtained from the corresponding region of Bovans brown legs that produce scales instead of feathers (Fig. 2b—Supplementary Information). To further enrich for genes associated with flight feather bud development, we also compared RNA-seq datasets from the posterior regions of Bovans brown legs and Pekin bantam legs (Fig. 2c—Supplementary Information). Pekin bantams develop feathered legs that have flight feathers along their posterior margins, whereas most chicken breeds including Bovans browns produce only scales ^26^.

**Figure 2.**
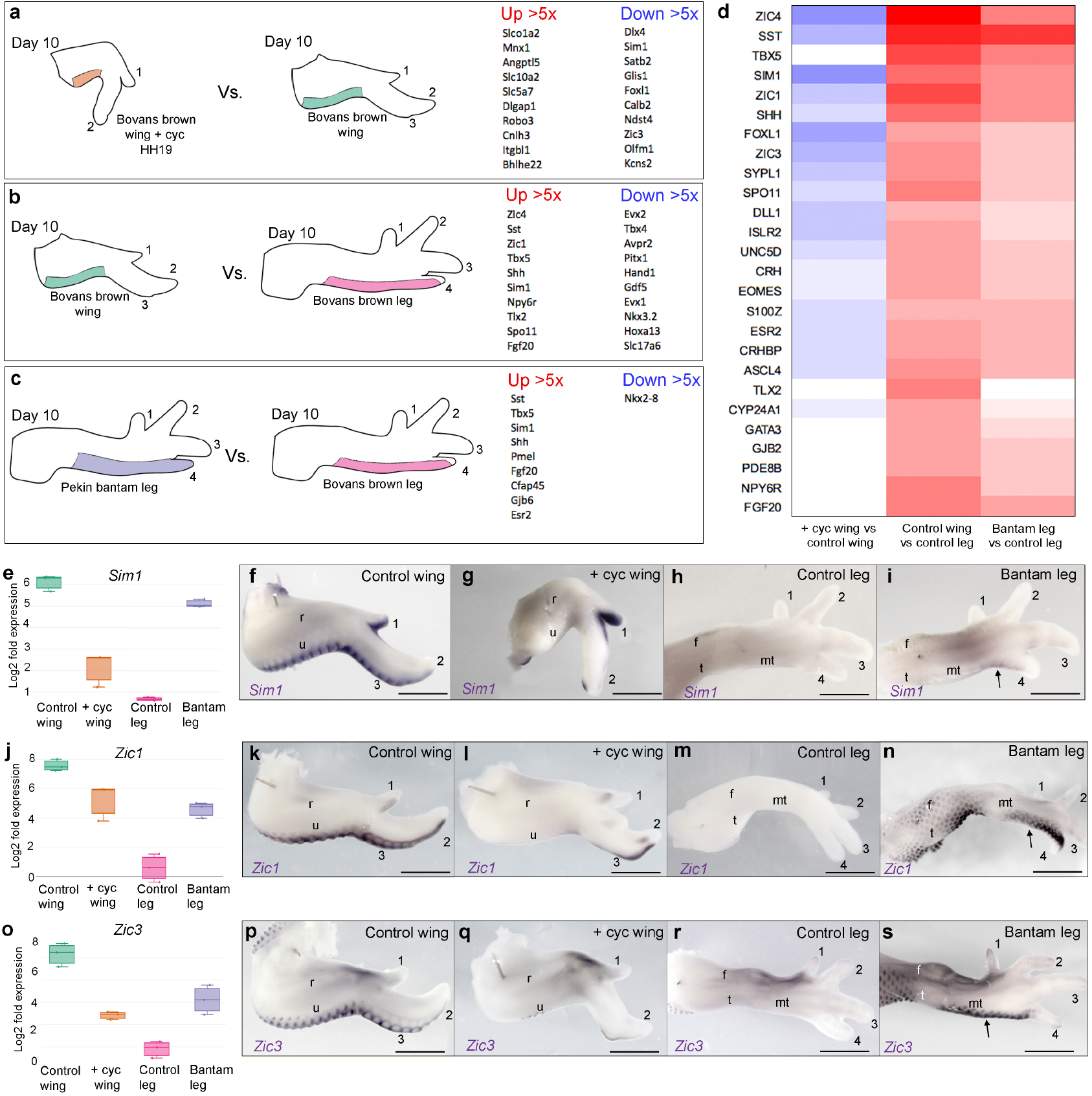
RNA-sequencing reveals targets of Shh signalling expressed in flight feather-forming regions. Schematics showing regions of day 10 limbs that were used to make RNA and the pairwise contrasts made: HH19 cyclopamine-treated Bovans brown wings vs. Bovans brown wings (**a**) HH19 Bovans brown wings vs. Bovans brown legs (**b**) HH19 Pekin bantam legs vs. Bovans brown wings (**c**)—top ten up-and down-regulated genes at >5-fold change are shown for each contrast (*p*-value of 0.005). Cluster of genes down-regulated in wings by earlier Shh signalling inhibition, up-regulated in wings vs. legs and up-regulated in Pekin bantam legs vs. Bovans brown legs (**d**—*p*-value of 0.005 and >2-fold change in at least one contrast; red—up-regulated, blue—down-regulated). Box-plots showing expression levels of *Sim1* (**e**), *Zic1* (**j**), and *Zic3* (**o**) as normalised log_2_ values of RNA sequencing read-count intensities. Expression of *Sim1* (*n*=22/22—**f**), *Zic1* (*n*=4/4—**k**), and *Zic3* (*n*=4/4—**p**) in flight feather-forming regions of day 10 wings—note *Sim1* expression in digit 1. Down-regulation of *Sim1* (*n*=12/14—**g**), *Zic1* (*n*=2/2—**l**), and *Zic3* (*n*=2/2—**q**) in forewing regions following Shh signalling inhibition. Undetectable/weak expression of *Sim1* (*n*=2/2—**h**), *Zic1* (*n*=2/2—**m**), and *Zic3* (**r**) in Bovans brown legs (t—tibia, f—fibula, mt—metatarsal). Up-regulation of *Sim1* (*n*=2/2—**i**), *Zic1* (*n*=2/2—**n**), and *Zic3* (*n*=2/2—**s**) along posterior margins of Pekin bantam legs. Scale bars: 1 mm

We performed a hierarchical clustering analysis to identify genes that behave similarly (genes included are expressed at a >2-fold difference between at least one contrast—*p*-value 0.005). This produced four clusters (Supplementary Information), and we focussed on cluster four that comprises twenty-six known genes including *Sim1* and *Zic1* (Figs. 2d, e and j—note, blue is down-regulated; red is up-regulated). As reported previously, *Sim1* (Fig. 2f) and *Zic1* Fig. 2k) are expressed in the flight feather-forming regions of the chick wing ^9 23^, although *Zic1* is only weakly expressed along the posterior margin of digit 1 (Fig. 2k). However, both *Sim1* (Fig. 2g) and *Zic1* (Fig. 2l) are undetectable along most of the posterior margin of the ulna and digit 2 of day 10 wings treated with cyclopamine at HH19, although *Sim1* is still observed in digit 1 (Fig. 2g). In addition, although both *Sim1* (Fig. 2h) and *Zic1* (Fig. 2m) are undetectable along the posterior margin of Bovans brown legs, they are expressed in equivalent regions of Pekin bantam legs (arrows—Figs. 2l and n). *Zic3* is also present in this cluster (Figs. 2d and o) and shares a very similar expression pattern with *Zic1* in normal wing development (compare Figs. 2k and p). In addition, as found with both *Sim1* and *Zic1*, the inhibition of Shh signalling reduces the expression of *Zic3* in forewing regions (Fig. 2q). Furthermore, compared with Bovans brown legs (Fig. 2r), *Zic3* is also strongly expressed along the posterior margins of Pekin bantam legs (Fig. 2s). Interestingly, this cluster also contains the *Tbx5* gene (Fig. 2d), which is implicated in feather formation in the legs of several bird species ^27^. These results provide evidence for a potential gene regulatory network operating downstream of Shh signalling in flight feather bud development.

### Shh signalling controls the spatial pattern *Sim1* expression

*Sim1* appears to be the clearest marker of the flight feather-forming regions of the chick wing from day 8 to day 13 Supplementary Figure 2—compare to *Bmp7* expression in all feather buds—Supplementary Figure 1). In order to precisely determine the temporal requirement for Shh signalling in specifying the later pattern of *Sim1* expression, we applied cyclopamine at a series of stages. Application at HH18 causes loss of *Sim1* expression along the posterior margin of the ulna and digit 2, and reduces expression in digit 1 (Fig. 3a). Progressively later treatments at HH19 cause loss of *Sim1* expression in the ulna and the proximal part of digit 2 (Fig. 3b), and at HH21, in digit 3 only (Fig. 3c). In addition, although *Shh* is expressed until HH28 12, treatment with cyclopamine after HH22/23 does not affect the pattern of *Sim1* expression (Fig. 3d). These findings reveal that Shh signalling from the polarising region between HH18 and HH22 specifies the later pattern of *Sim1* expression in a defined spatial sequence. Thus, in reference to the classical positional information of digit patterning ^16^, a short exposure of Shh signalling (low concentration) is sufficient for expression of *Sim1* in digit 1, and progressively longer exposures (higher concentrations), for expression in the distal part of digit 2, the ulna, and then digit 3.

**Figure 3.**
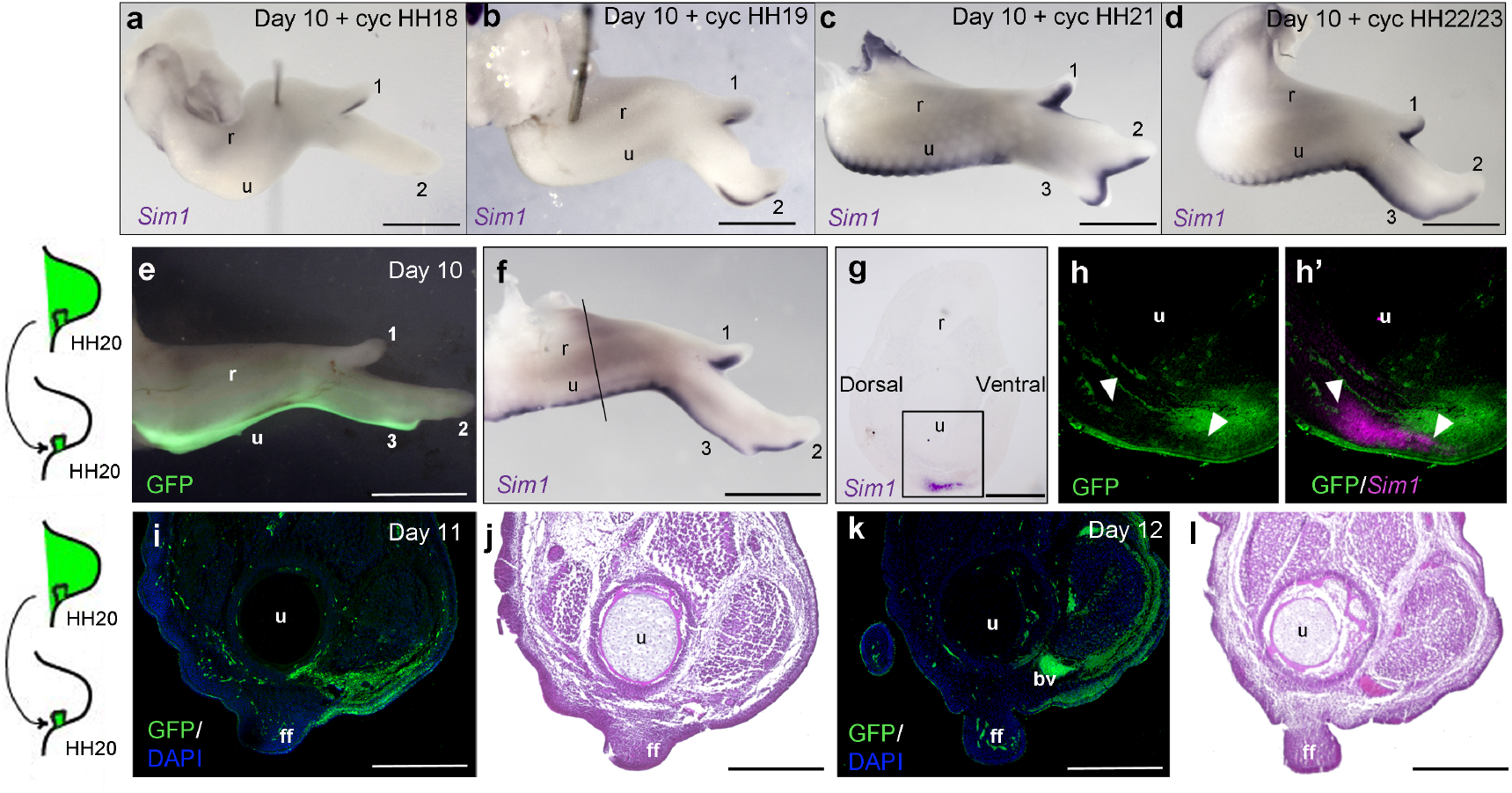
Shh signalling specifies the spatiotemporal pattern of *Sim1* expression. Application of cyclopamine at HH18 reduces *Sim1* expression in digit 1 at day 10 and causes loss of expression in the ulna and digit 2 (*n*=3/3—**a**), at HH19—loss of *Sim1* expression in the ulna and the proximal region of digit 2 (*n*=12/14—**b**), at HH21—loss of *Sim1* expression in digit 3 (*n*=10/15—**c**), and after HH22, the pattern of *Sim1* expression is unaffected (*n*=5/5—**d**). GFP-expressing polarising regions transplanted in place of normal polarising regions at HH20 contribute to posterior soft tissues of day 10 wings (*n*=2/2—**e**—r-radius, u-ulna, digits 1, 2, and 3) and resemble *Sim1* expression along posterior margin of the ulna and digit 2 (**f**). Transverse section through forewing shown in e reveals adjacent expression of GFP and *Sim1* in dermis (*n*=2/2—**g**, **h, h’**—note GFP protein and *Sim1* mRNA detected on same section). Transverse section through day 11 wing (*n*=2/2—**i**, experiment performed as in a) and day 12 wing (*n*=2/2—**k**) showing GFP expression ventral to emerging flight feather (ff, blood vessel—bv and blue shows DAPI staining). H&E staining on serial sections to i and k shows tissue anatomy (**j**, **l**). Scale bars: a-f—1 mm; g-h—150 μm; i-l—75 μm

The pattern of *Sim1* expression along the posterior margin of the wing superficially resembles polarising region fate maps ^14^. Therefore, to examine this lineage relationship, we replaced normal HH20 polarising regions with HH20 GFP-expressing polarising regions, and then analysed the expression of *Sim1* seven days later (Figs. 3e and f). Transverse sections show polarising region-derived GFP-expressing dermal cells lying immediately ventral to both *Sim1*-expressing cells in forewings at day 10 (Figs. 3g and h) and emerging flight feathers at days 11 and 12 (Figs. 3i-l). Taken together, these results demonstrate that Shh signalling by the polarising region is required for later *Sim1* expression and flight feather bud formation in a defined spatial and temporal sequence.

### Shh signalling is required for flight feather formation in hatched chicks

During the stages leading up to hatching (days 18-21 of incubation), the first generation of flight feathers is replaced by the second generation of mature flight feathers ^1 2^. Therefore, to determine if polarising region-derived Shh signalling is required for the formation of mature flight feathers, we allowed chicks treated with cyclopamine at HH19 to hatch on day 21 of incubation. In untreated wings, both well-formed flight feathers and dorsal major covert feathers can be observed extending away from the posterior margin of the wing (Figs. 4a and a’—note natal down trimmed back). H&E staining on a transverse section through the forewing region shows the ligament connecting the flight feather to the ulna (Figs. 4b and b’).

**Figure 4.**
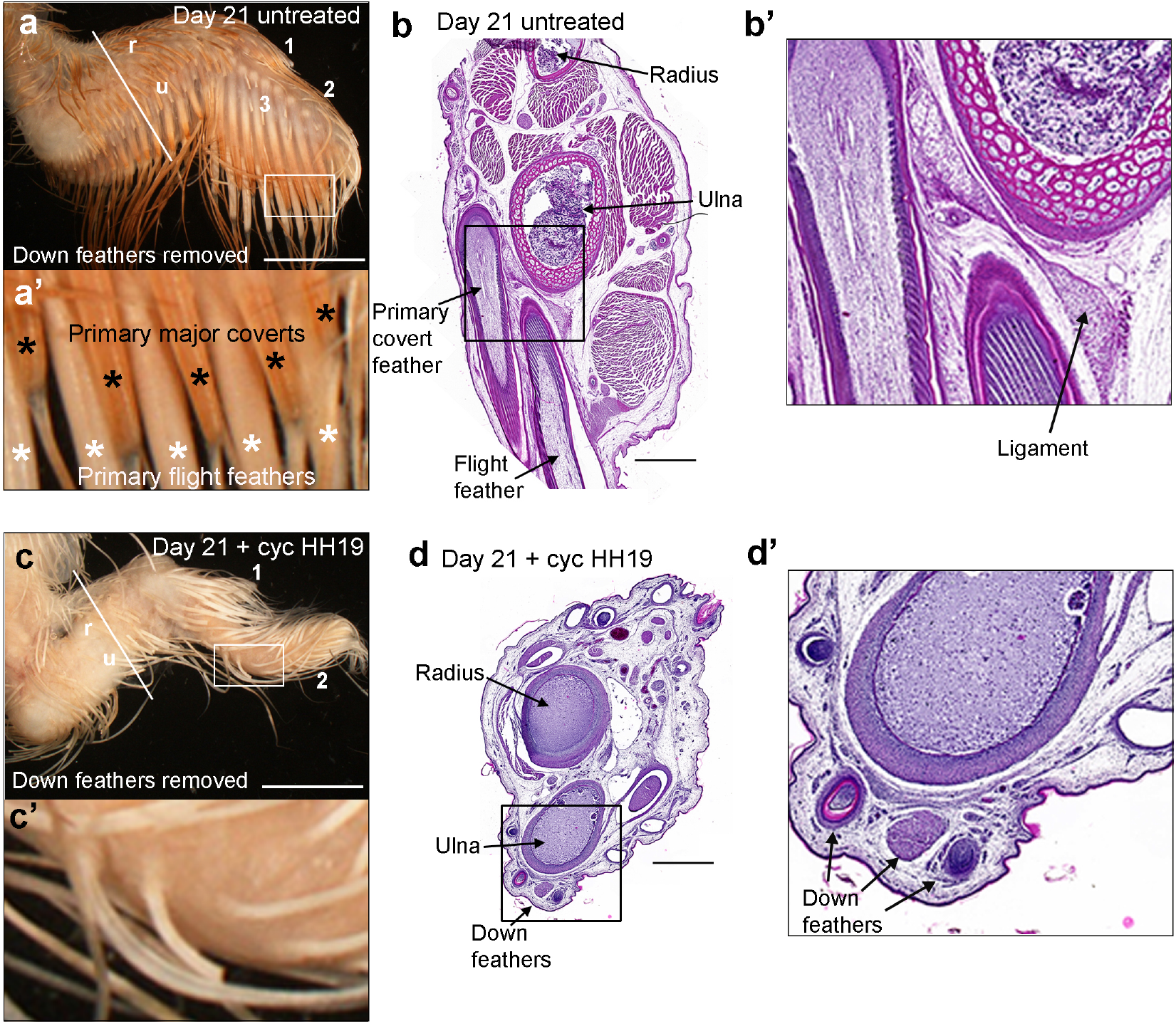
Shh signalling is required for flight feather formation in the wings of hatched chicks. Example of an untreated chicken wing at hatching (incubation day 21) showing the normal pattern of primary flight feathers and primary major covert feathers (**a**, **a**’ *n*=13/16). H&E staining on a transverse section through the wing in (a) show ligaments connecting the flight feathers to the ulna (**b**, **b**). Example of a HH19 cyclopamine-treated wing at hatching showing loss of both primary flight feathers and primary major covert feathers (**c**, **c’** *n*=13/16). H&E staining on a transverse section through the wing in (c) shows that down feathers are still present at the posterior margin of the wing where flight feathers would normally develop (**d**, **d**’). Scale bars: a, c—8 mm; b, d—1 mm.

However, in chicks treated with cyclopamine at HH19, both flight feathers and dorsal major covert feathers fail to develop at hatching (Figs. 4c and c’—note natal down trimmed back - Supplementary Table 1). It is worth noting that natal down feathers still form along the posterior margin of the wing in regions where flight feathers would normally form, thus showing that inhibition of Shh signalling does not result in the general inability to initiate feather development (Figs. 4d and d’). Additionally, there is a precise time-window during which Shh signalling is required for later flight feather formation, since flight feathers often form in distal regions of wings treated with cyclopamine at HH20/21 (Supplementary Figure 3 and Supplementary Table 1). These are the exact locations where restricted expression of *Sim1* is often observed following the earlier inhibition of Shh signalling (Figs. 3a and b). In addition, the flight feather pattern is usually normal following cyclopamine treatment at HH22/23 (Supplementary Figure 3 and Supplementary Table 1), just as the *Sim1* expression pattern in also normal (Fig. 3d). Therefore, Shh signalling by the polarising region specifies the spatial pattern of *Sim1* expression and flight feather formation in the same temporal sequence.

### Shh signalling is required for flight feather formation in mature birds

Chicks that were treated with cyclopamine at HH19 did not survive beyond hatching, which prevented the study of their mature wing plumage. Therefore, since our analyses of hatched chickens shows that dorsal major coverts—which are closely associated with developing flight feathers—are also absent, this raised the possibility that Shh inhibition could affect the later development of other feathers that had not yet replaced the natal down.

To analyse feather development in mature bird wings, we treated embryos at HH19/20 with cyclopamine and then grafted their right-hand wing buds in place of the right-hand wing buds of untreated embryos (Fig. 5a – see control experiment showing that the grafting procedure does not affect feather development – Supplementary Figure 4). This procedure enabled chicks to survive beyond hatching (Fig. 5b), and they displayed the same patterns of flight feather loss as hatched chicks that were systemically treated with cyclopamine as embryos (Supplementary Tables 1 and 2). Several birds were allowed to progress to later stages of development and their patterns of flight feather loss remained the same as at hatching (Supplementary Table 2). Thus, one such example of a postnatal day 22 bird shows that the flight feather pattern is normal in its untreated left-hand wing, but that there is a loss of distal primary flight feathers in its cyclopamine-treated right-hand wing (arrow, Fig. 5c). This bird was allowed to survive until postnatal day 66, so that its adult feather pattern could be studied in more detail (Figs. 5d-g). Manual examination of its untreated left-hand wing reveals that the natal down has been replaced by defined rows of mature feathers (dorsal view Fig. 5d, ventral view Fig. 5f). Thus, eighteen secondary flight feathers develop from its ulnar region (green asterisks Figs. 5d and f, note - not all feathers can be seen because they overlap) and ten primary feathers develop from its digital region (eight primaries from digit 3—orange asterisks, and two from digit 2—blue asterisks, Figs. 5d and f, note, one feather was broken). Three alular flight feathers extending from digit 1 are also present (red asterisks Figs. 5d and f). Lying immediately anterior to the primary and secondary flight feathers are rows of dorsal and ventral major covert feathers (purple asterisks Figs. 5d and f), and above these, median covert feathers are easy to distinguish on the dorsal side of the wing (light blue asterisks Fig. 5d). The remaining feathers that are present in more-anterior regions of the wing can be classified as marginal covert feathers (Figs. 5d and f ^1^). However, in the contralateral cyclopamine-treated wing of this bird, eight primary flight feathers are absent along the posterior margin of digit 3 and the dorsal major coverts overlying them are also absent (Figs. 5e and g). In addition, two bunched primary flight feathers—which are much smaller than the equivalent ones in its control wing—extend from the margin of digit 2 at the distal tip of the wing (blue asterisks - Figs. 5e and g), and overlying them, are dorsal major covert feathers (purple asterisks - Fig. 5e). The remaining feathers are of the same identities as those present in its untreated wing, although some of the secondary flight feathers are shorter (green asterisks – Figs. 5e and g). In addition, the development of ventral major covert feathers is unaffected by cyclopamine treatment (Fig. 5g). Interestingly, the pattern of feather loss in the wing of this bird is consistent with the pattern of *Sim1* expression in the wings of embryos that were treated with cyclopamine at HH19 (Fig. 3b). These results demonstrate that the inhibition of Shh signalling in the embryo causes the selective loss of mature flight feathers and their overlying dorsal major coverts in a defined spatial and temporal sequence.

**Figure 5.**
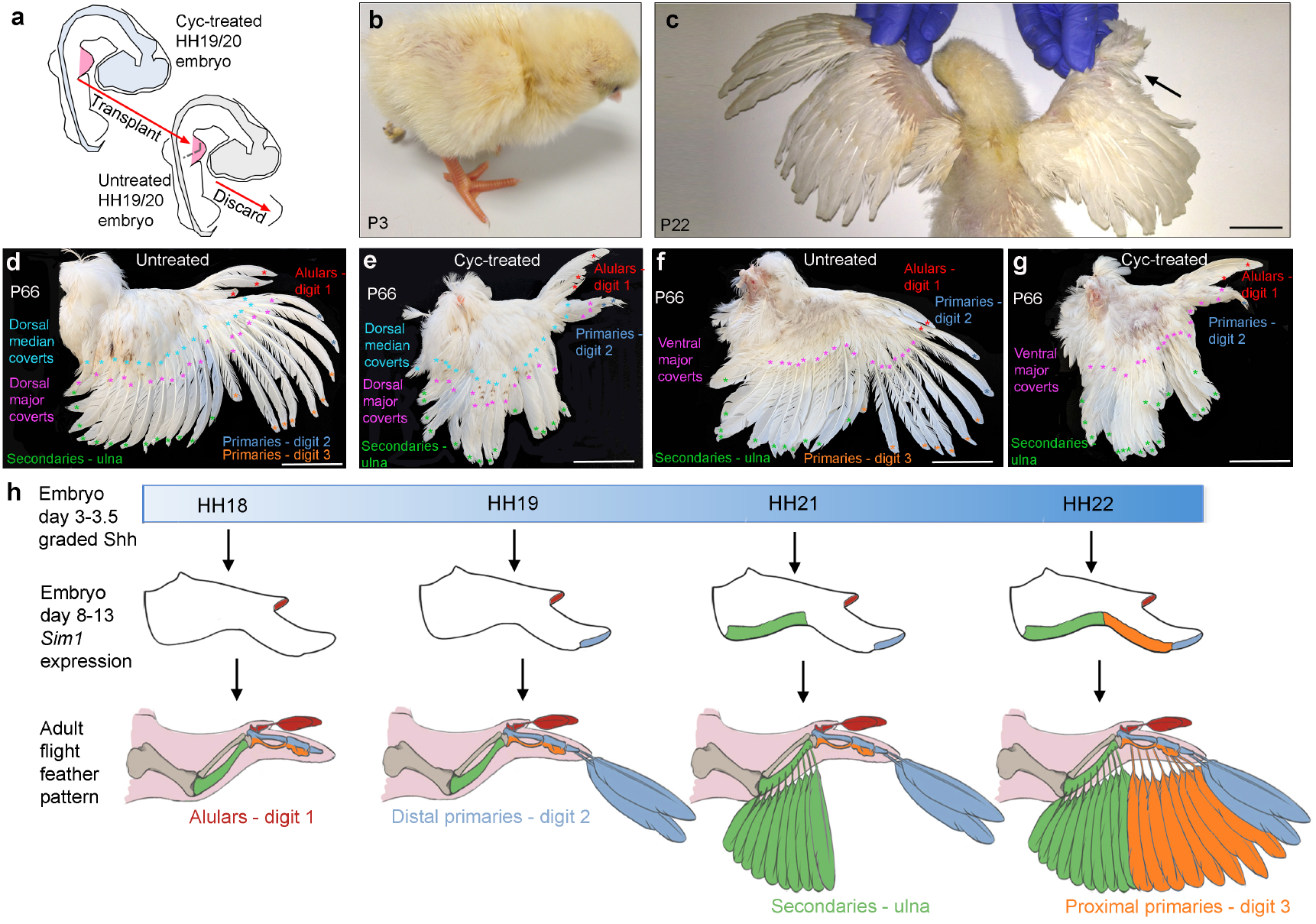
Shh signalling is required for flight feather formation in the wings of mature birds. Experimental procedure in which a wing bud from a HH19/HH20 cyclopamine-treated embryo was grafted in place of a wing bud of an untreated embryo (**a**). Example of a hatched chick (p3 – postnatal day 3 - **b**) that underwent the procedure described in (a). Same chicken shown in (b) at p22 – note, absence of primary flight feathers in distal regions of the cyclopamine-treated wing (arrow-**c**). Dorsal views of untreated (**d**) and cyclopamine-treated wings at p66 (**e**, same chicken in b, c) showing alular flight feathers (red asterisks, digit 1), distal primary flight feathers (blue asterisks, digit 2), proximal primary flight feathers (orange asterisks, digit 3), secondary flight feathers (green asterisks, ulna), dorsal major covert feathers (purple asterisks) and dorsal median covert feathers (light blue asterisks). Note absence of proximal primary flight feathers and overlying primary major covert feathers in cyclopamine-treated wing (**e**). Ventral views of untreated (**f**) and cyclopamine-treated wings (**g**) at p66 showing alular flight feathers (red asterisks, digit 1), distal primary flight feathers (blue asterisks, digit 2), proximal primary flight feathers (orange asterisks, digit 3), secondary flight feathers (green asterisks, ulna) and ventral major covert feathers (purple asterisks). Note absence of proximal primary flight feathers in cyclopamine-treated wing (**g**). Graded Shh signalling between HH18 and HH22 (blue shading—**h)**specifies the spatiotemporal pattern of *Sim1* expression and adult flight feathers in the order that the pattern of skeletal elements is also specified across the antero-posterior axis. Scale bars: c—3 cm; d-g—5 cm

## Discussion

### Embryonic Shh signalling is required for flight feather formation

We have revealed that Shh signalling by the embryonic chick wing polarising region is required for specifying the adult pattern of flight feathers and their associated dorsal major covert feathers. This process is independent of the later role that Shh signalling fulfils in feather morphogenesis ^19–21^. Thus, the transient ablation of Shh signalling between HH18 and HH22, but not later, causes the loss of bilaterally asymmetric flight feathers that make ligamentous connections to the skeleton, and also the loss of molecular markers associated with the flight feather forming regions of wing. However, the Shh signalling pathway (*Ptch1*) is still active during later feather bud morphogenesis, thereby showing that this general process is unaffected.

Detailed fate mapping experiments have shown that chick wing bud cells contribute to the development of distal structures when Shh signalling by the polarising region is transiently blocked ^15 14^. Taken together with the finding that apoptosis is also suppressed in the posterior part of the wing bud 18, this provides evidence that the loss of both dorsal major covert and flight feathers is not caused by the selective loss of cells. We also revealed that the dorso-ventral boundary, which is important for flight feather development ^22^, remains intact following the inhibition of Shh signalling. In addition, we demonstrated that feather buds, which produce other feather types, still form along the posterior border of the ulna and digit skeleton. Therefore, our data provides molecular insights into classical tissue recombination experiments, which showed that feather position and identity are determined by signals acting at around HH20 ^10,11^.

### A positional information gradient of Shh specifies flight feather identity

Our findings can be explained by the classical positional information model of antero-posterior patterning, in which Shh signalling specifies limb bud cells with a positional value, which when interpreted at a later stage of development, allows them to differentiate into the appropriate structure ^16,17^. Thus, the temporal requirement for Shh signalling in specifying the anterior to posterior pattern of *Sim1* expression and flight feathers closely follows that for specifying the anterior to posterior pattern of digits ^13 14 16^ (Fig. 5h): digit 1 and alular flight feathers are specified first by a low concentration/short duration of Shh at HH18, and then increasing concentrations of Shh over time specify the other skeletal elements and flight feathers in the order; digit 2 and distal primaries at HH19; the ulna and secondaries at HH21, and digit 3 and proximal primaries at HH22 (Fig. 5h). It is of note that the pattern of *Sim1* expression in the wings of embryos treated at HH19, precisely matches the pattern of flight feathers present both at hatching and in mature bird wings (Fig. 5h). Therefore, eight proximal primaries—which normally form along the border of digit 3—are absent; yet two distal primaries are present along the border of digit 2 (Fig. 5h). This pattern of flight feather loss is also accompanied by the loss of the associated overlying row of dorsal major covert feathers. However, the inhibition of Shh signalling does not affect the development of the remaining feathers in the wing, thereby implying that their identities are specified by other signals. Our findings therefore indicate that flight feathers and dorsal major covert feathers have similar developmental programmes, the study of which could warrant further investigation. Taken together, these observations reveal that the evolution of the flight feather programme involved the co-option of the pre-existing positional information gradient of Shh signalling used in forewing/digit patterning (Fig. 5h).

### Interpretation of the Shh gradient into flight feather identity

The interpretation of positional information, in which cells memorise their positional value to give rise to appropriately patterned and positioned structures at a later stage of development, is generally an unknown process in developmental biology ^28^. Indeed, despite decades of research, the genes acting downstream of the positional information gradient of Shh signalling in the specification of digit identity remain largely unknown ^16^. However, our RNA sequencing experiments provide molecular insights into a putative gene regulatory network that operates downstream of Shh signalling in determining flight feather identity. These genes include *Sim1*, and notably, genes encoding three Zic transcription factors (*Zic1*, *3* and *4*). Interestingly, Zic transcription factors can bind to sites in promoters that are also recognised by the Gli family of transcription factors—the downstream effectors of Shh signalling ^29^. This mechanism could provide a positional memory mechanism in which polarising region-derived Shh signalling could remove Gli transcriptional repressors from the promoters of genes, thus making them accessible to Zic transcription factors at stages of flight feather bud development. Such directions could be the focus of future studies.

In conclusion, as flight feathers were one of the earliest known adaptations associated with the evolution of flight in theropod dinosaurs ^3–5^, our findings have significant implications for this extraordinary transition.

## Methods

### Chick husbandry and tissue grafting

Wild type and GFP-expressing Bovans brown chicken eggs were incubated and staged according to Hamburger Hamilton ^30^. Day 3 of incubation is HH18, day 4 - HH21, day 5 - HH24, day 6 – HH27, day 7 - HH29, day 8 – HH30, day 10 - HH36, day 11 – HH37, day 12 – HH38, day 13 – HH39, day 14 – HH40 – day 15 – HH41 and hatching at day 21 is HH46. All experiments involving the use of hatching chicken embryos in this work were conducted in accordance with the EU animal experiment guidelines and reviewed and approved by the Bioethics Committee of the University of Cantabria (PI-20-17).

#### Polarising region and wing bud grafts

Embryos were dissected in DMEM and polarising regions or wing buds removed using fine tungsten needles, grafted to the appropriate location of stage-matched host limb buds and held in place with 25 μm platinum pins. Polarising region tissue was removed in reference to patterns of *Shh* expression.

#### H&E staining

12 μm transverse sections of paraffin embedded forewings were mounted on glass slides. Slides were washed twice in xylene for 5 mins followed by rehydration through an ethanol series (2x 100%, 95%, 70%) and were washed in H_2_O. Slides were stained for 2 mins in Harris haematoxylin followed by differentiation in 0.3% acid alcohol. Blueing was achieved in Scott’s tap water and slides were rinsed in H_2_O before staining in eosin for 5 mins. Slides were rinsed in H_2_O and dehydrated through ethanol series (70%, 95%, 100%). Dehydrated slides were cleared of remaining wax with xylene before mounting.

#### Shh signalling inhibition

Cyclopamine (Sigma) suspended in control carrier (45% 2-hydropropyl-β-cyclodextrin in PBS, Sigma, to a concentration of 1mg/ml) and 4 μl pipetted directly onto embryos over the limb bud, after removal of vitelline membranes. In all cases, untreated wings were treated with 2-hydropropyl-β-cyclodextrin only. Digit identities were determined by visualising phalange position under illumination.

#### Whole mount RNA in situ hybridisation

Embryos were fixed in 4% PFA overnight at 4°C, dehydrated in methanol overnight at −20°C, rehydrated through a methanol/PBS series, washed in PBS, then treated with proteinase K for 20 mins (10μg/ml^−1^), washed in PBS, fixed for 30 mins in 4% PFA at room temperature and then pre-hybridised at 65°C for 2 h (50% formamide/50% 2x SSC). 1μg of antisense DIG-labelled (Roche) mRNA probes were added in 1 ml of hybridisation buffer (50% formamide/50% 2x SSC) at 65°C overnight. Embryos were washed twice in hybridisation buffer, twice in 50:50 hybridisation buffer and MAB buffer, and then twice in MAB buffer, before being transferred to blocking buffer (2% blocking reagent 20% lamb serum in MAB buffer) for 2 h at room temperature. Embryos were transferred to blocking buffer containing anti-digoxigenin antibody (Roche 1:2000) at 4°C overnight, then washed in MAB buffer overnight before being transferred to NTM buffer containing NBT/BCIP and mRNA distribution visualised using a LeicaMZ16F microscope.

### Double RNA in situ hybridisation/immunohistochemistry

Wholemount RNA in situ hybridisation was performed as above. Embryos were fixed for 20 mins at room temperature in 4% PFA, washed 2x in PBT for 10 mins and then dehydrated through an ethanol series (10 mins each wash in PBT) to 100% EtOH and stored at −20°C overnight. Embryos were cleared in xylene until light was visible through the tissue (approx. 2-10 mins). Embryos were processed through a series of 30 min wax changes at 60°C (25%, 25%, 50%, 75%, 100%, 100%) and then left in the oven overnight. Limbs were embedded in wax and allowed to set for 4-6 h before being sectioned using a microtome and the sections were floated on a slide rack overnight at 52°C. Slides were washed in xylene for 5 mins (2x) in a Coplin jar then rehydrated through an ethanol series (2x 5 mins washes each) to H_2_0 and then washed twice in PBT. Slides were blocked horizontally for 1 hour in 3% HINGS in PBT and incubated in primary antibody (anti-chick GFP at 1:100) in blocking solution overnight at 4°C or for 4 h at room temperature. Slides were washed in a Coplin jar (3x 15-30 mins) and then incubated in secondary antibody goat anti-chicken conjugated to Alexa 488 at 1:500) in blocking solution in the dark. Slides were rinsed 4-5 times in the dark in PBS and mount with Fluoroshield (with DAPI).

### RNA sequencing analyses and clustering

Tissue used for making RNA was manually dissected using fine forceps. Three replicate experiments were performed from each condition and the tissue was pooled before the RNA was extracted using Trizol reagent (Gibco). Sequencing libraries were prepared using Illumina TruSeq library preparation kit. Samples were sequenced on a HiSeq 2000 (Paired end readings of 50 bp - Instrument: ST300). Reads were aligned to the chicken genome, assembly Gallus_gallus-5.0, using STAR aligner.

The raw data has been deposited in array express (https://www.ebi.ac.uk/arrayexpress/experiments/E-MTAB-7520). A total of 12 samples (3 replicates for each condition) were QC analysed using automatic outlier detection. This was done by manually inspecting the density plot, boxplots, PCA plots, correlation heatmap and distance plot, as well as using several automatic outlier tests, namely distance, Kolmogorov-Smirnov, correlation and Hoeffding’s D (all samples passed QC). The count data for the samples were normalised using trimmed mean of M-values normalisation and transformed with voom, resulting in log_2_-counts per million with associated precision weights.

Genes were clustered using the clValid R package based on their log_2_ fold changes. The Dunn Index was selected as the preferred cluster validation measure. Three clustering methods (hierarchical, k-means and PAM) were tested for two up to 20 clusters and the clustering analysis was performed on 906 unique genes that were differentially expressed in these contrasts at the significance threshold of FDR-adjusted p-value < 0.005 and fold change ≥ 2. Using k-means the clustering of the 906 genes resulted in four groups.

## Supporting information

RNA seq data

## Acknowledgements

We acknowledge Marysia Placzek, Cheryll Tickle and Jasmina Tarpy for critical reading, Max Bylesjo, Laura Bennett and Dan Halligan (FIOS genomics) for bioinformatics and Adrian Sherman (Roslin Institute) for transgenic chicks and James Briscoe for the *Sim1* plasmid. MT thanks the Wellcome Trust for funding (202756/Z/16/Z) and MR the Spanish Ministerio de Ministerio de Ciencia, Innovación y Universidades (BFU2017-88265-P).

## Author Contributions

LB performed most analyses of day 8 to day 15 chicks and edited the paper; CA performed most analyses of day 15 to day 21 chicks; CM performed RNA-seq validation and contributed to section in situs/immunos and H&E staining; CR contributed to RNA preparation for RNA-seq and to section in situs/immunos. MR contributed to the analyses of day 15 – day 21 chicks, did all of the post-hatching work, transplant experiments and edited the paper; MT devised the study, performed pilot experiments and wrote the paper.

## Author Information

- The authors declare no competing interests

---

**Supplementary Figure 1.**
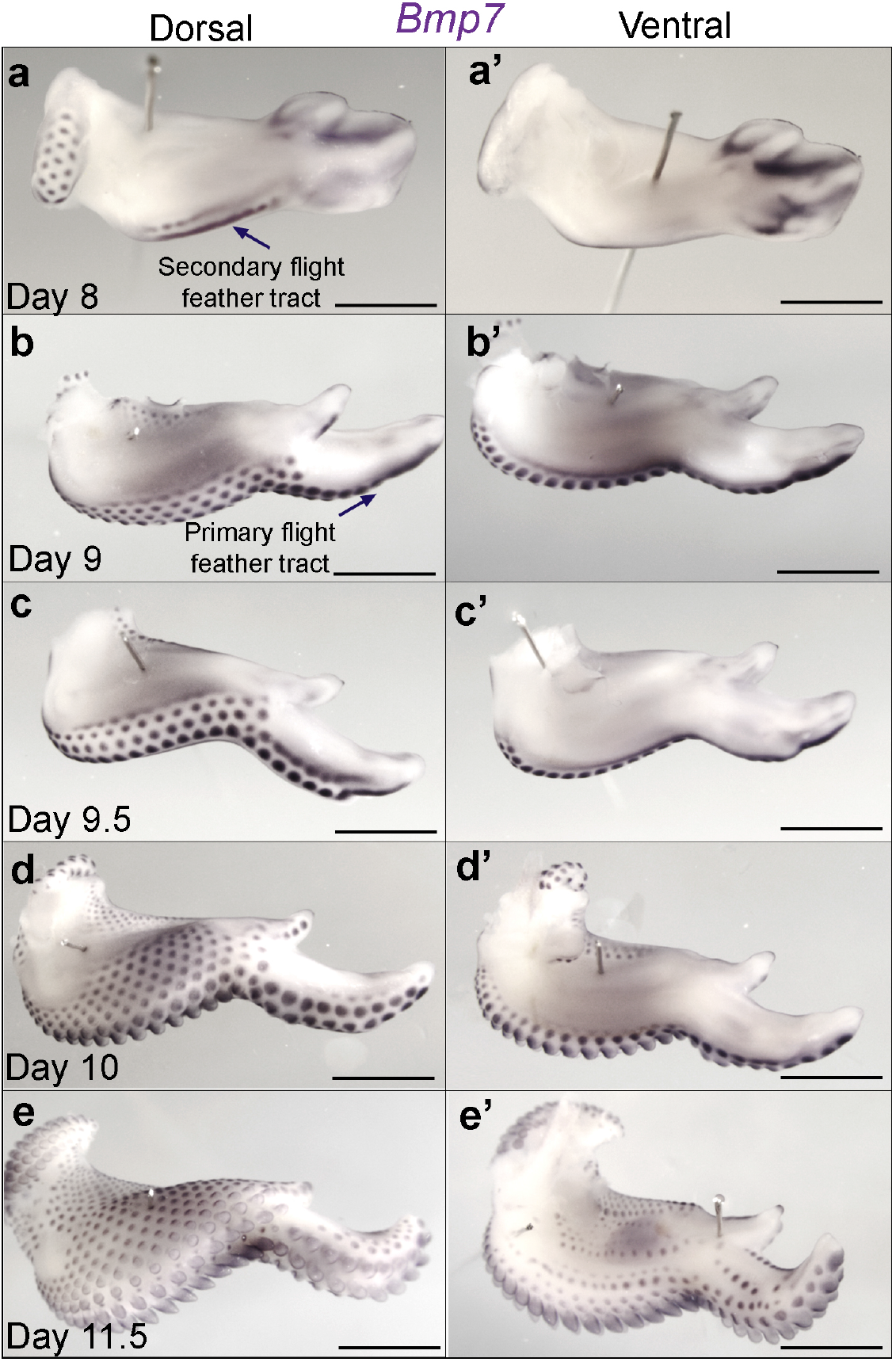
Expression of *Bmp7* during chick wing feather development. *Bmp7* is expressed in all feather buds. Note, dorsal and ventral views are shown and tracts of feather buds form in a posterior to anterior sequence over time (**a-e**)—the first tract to form in the forewing is the secondary flight feather tract at day 8 (**a**) — the first tract to form in the hand-plate is the primary flight feather tract at day 9 (**b**). Scale bars: 1 mm

**Supplementary Figure 2.**
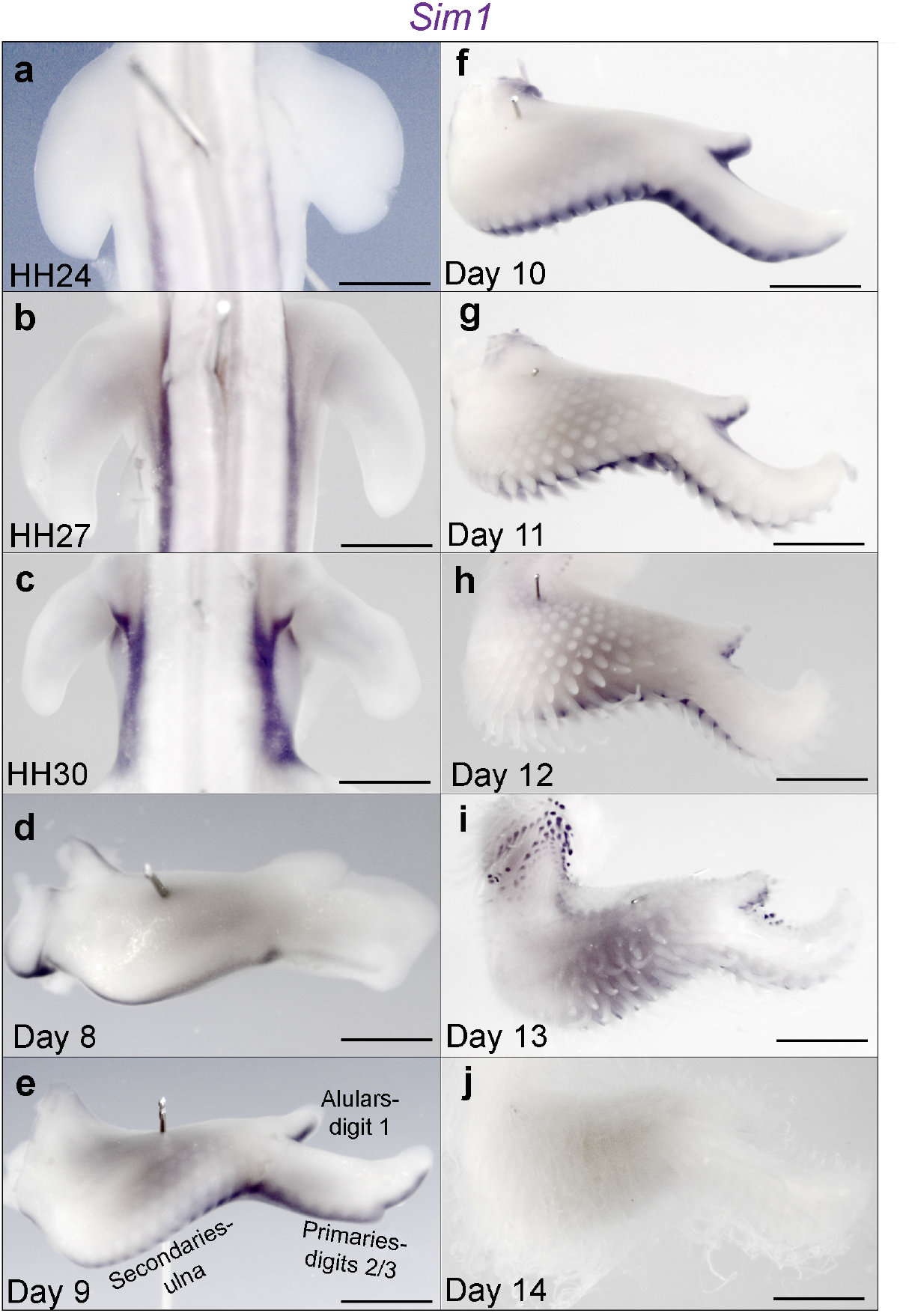
Expression of *Sim1* during chick wing development. *Sim1* is expressed in regions of flight feather development between day 8 and day 14 (**a**-**j**). Scale bars: 1 mm

**Supplementary Figure 3.**
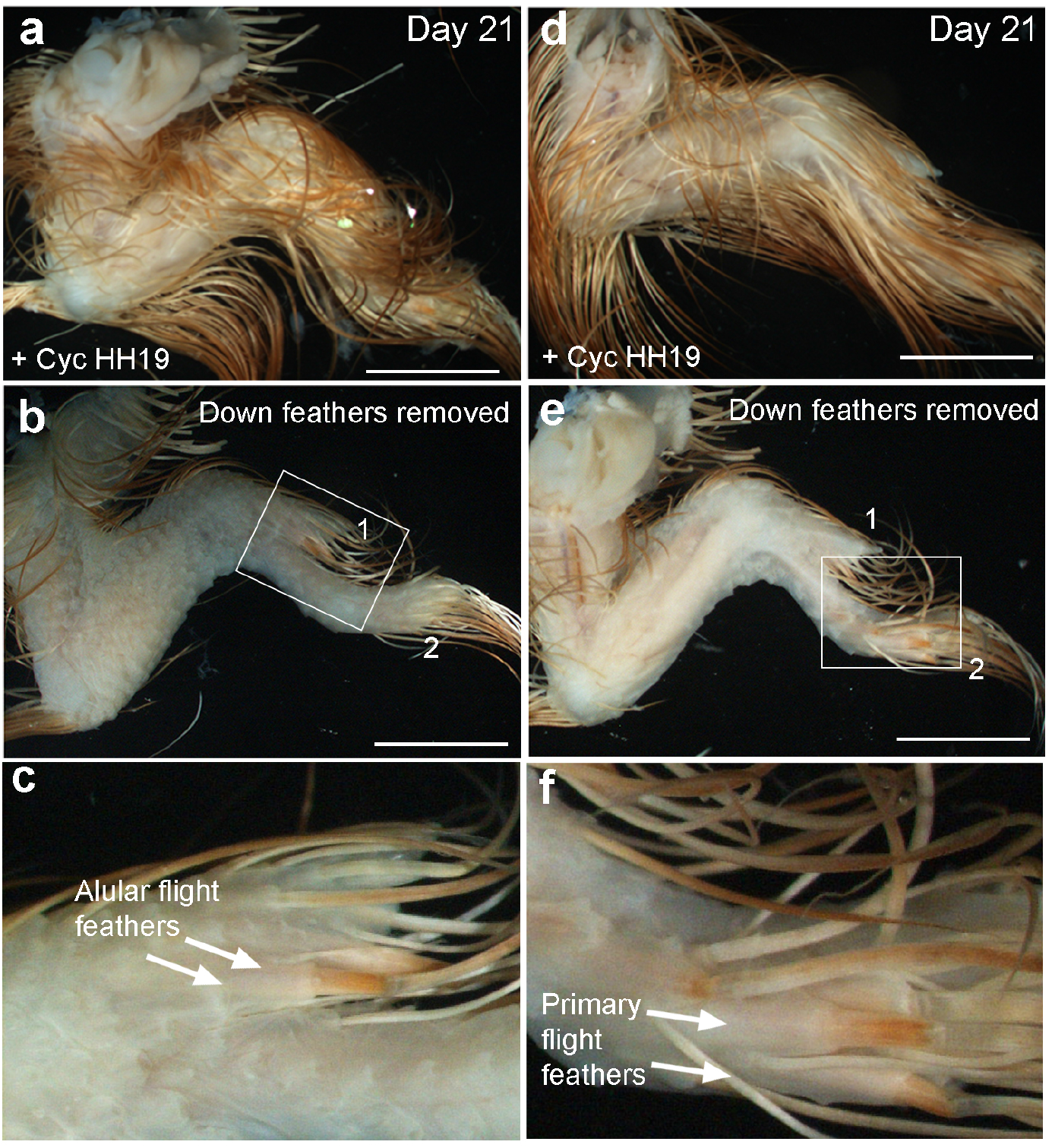
Flight feather development in cyclopamine-treated wings at hatching. Two examples of HH19 cyclopamine-treated wings at hatching (**a**, **d**) in which flight feathers can be only observed in distal regions of digit 1 (**b**, **c**—alulars), and digit 2 (**e**, **f**—distal primaries)—See Supplementary Table 1. Scale bars: 8 mm

**Supplementary Figure 4.**
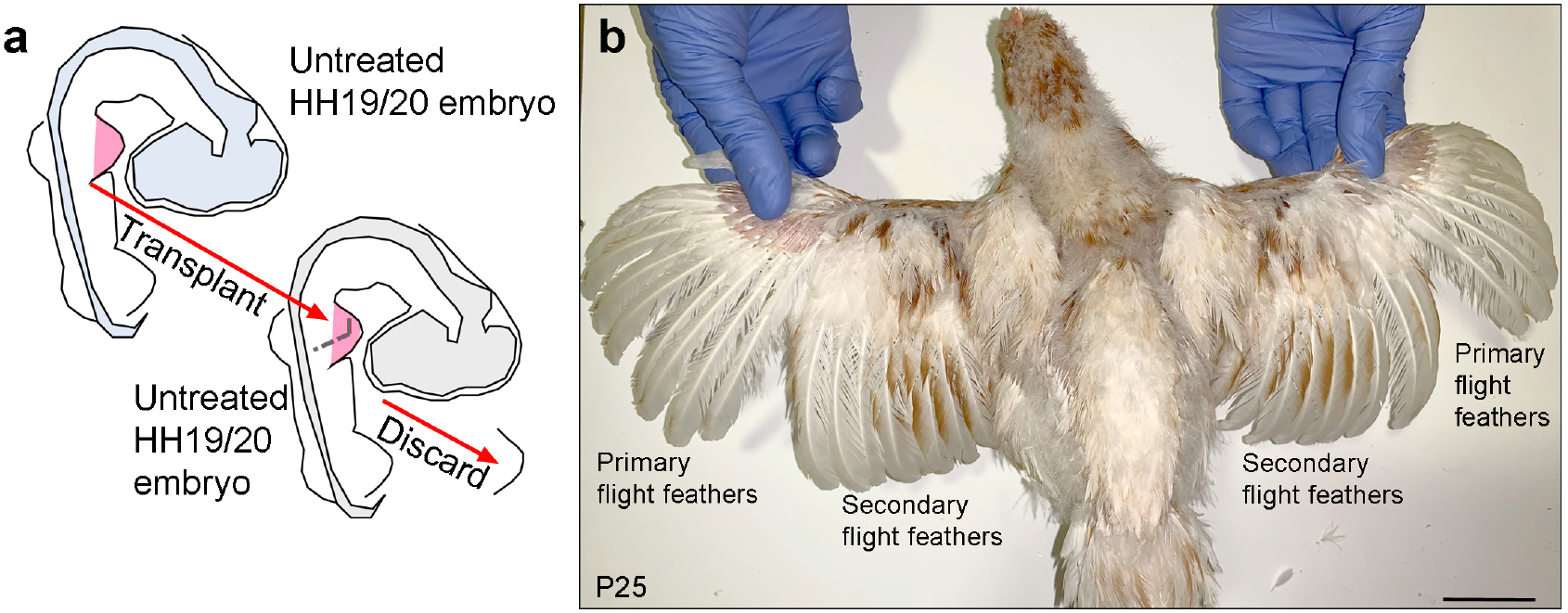
Normal mature feather development in transplanted wings. Schematic showing procedure in which an untreated HH19/20 wing bud is grafted in place of a stage-matched wing bud of another embryo (**a**). A chicken that underwent this grafting procedure showing normal development of feathers at p25 (**b**). Scale bar: 5 cm

**Supplementary Table 1.** effects of cyclopamine on feather pattern in day 21 chickens at hatching.

|  | Untreated<br>(n=10) | Cyc HH19<br>(n=16) | Cyc HH20/21<br>(n=19) | Cyc<br>HH22/23<br>(n=3) |
| --- | --- | --- | --- | --- |
| Normal flight feathers | 10 | 3 | 6 | 2 |
| Some rudimentary distal primary flight feathers and alulars. No secondaries | 0 | 11 | 2 | 1 |
| Some distal primaries and secondaries | 0 | 2 | 11 | 0 |

**Supplementary Table 2.** effects of cyclopamine on feather pattern on transplanted wings grafted to host normal chick embryos at hatching and in later development.

|  | Untreated<br>(n=3) | Cyc HH20<br>(n=3) | Cyc HH20/21<br>(n=3) |
| --- | --- | --- | --- |
| Normal flight feathers | 3 | 0 | 0 |
| Some rudimentary distal primary flight feathers and alulars. No secondaries | 0 | 2 | 1 |
| Some distal primaries and secondaries | 0 | 1 | 2 |

## References

1 Lucas, A. M. & Stettenheim, P. R. Avian Anatomy - Integument. Agriculture handbook 362 (1972).

2 Kondo, M. et al. Flight feather development: its early specialization during embryogenesis. Zoological Lett 4, 2, doi:10.1186/s40851-017-0085-4 (2018).

3 Turner, A. H., Makovicky, P. J. & Norell, M. A. Feather quill knobs in the dinosaur Velociraptor. Science 317, 1721, doi:10.1126/science.1145076 (2007).

4 Ortega, F., Escaso, F. & Sanz, J. L. A bizarre, humped Carcharodontosauria (Theropoda) from the lower cretaceous of Spain. Nature 467, 203–206, doi:10.1038/nature09181 (2010).

5 Godefroit, P. et al. Reduced plumage and flight ability of a new Jurassic paravian theropod from China. Nat Commun 4, 1394, doi:10.1038/ncomms2389 (2013).

6 Chen, C. F. et al. Development, regeneration, and evolution of feathers. Annu Rev Anim Biosci 3,169–195, doi:10.1146/annurev-animal-022513-114127 (2015).

7 McCrady, E. THE “PENGUIN” GUINEA FOWL: Absence of Flight Feathers Due to Hereditary Local Alopecia*. The Journal of Heredity 23,201–207 (1932).

8 Urrutia, M. S., Crawford, R. D & Classen, H. L. Dysplastic remiges, a genetic abnormality reducing feathering in the domestic fowl. The Journal of Heredity 74,101–104 (1983).

9 Seki, R. et al. Functional roles of Aves class-specific cis-regulatory elements on macroevolution of bird-specific features. Nat Commun 8, 14229, doi:10.1038/ncomms14229 (2017).

10 Cairns, J. M. & Saunders, J. R. The influence of embryonic mesoderm on the regional specification of epidermal derivatives in the chick. J. Exp Zoo 127,221–248 (1954).

11 Saunders, J. W., Jr. & Gasseling, M. T. Effects of reorienting the wing-bud apex in the chick embryo. J Exp Zool 142,553–569 (1959).

12 Riddle, R. D., Johnson, R. L., Laufer, E. & Tabin, C. Sonic hedgehog mediates the polarizing activity of the ZPA. Cell 75,1401–1416. (1993).

13 Yang, Y. et al. Relationship between dose, distance and time in Sonic Hedgehog-mediated regulation of anteroposterior polarity in the chick limb. Development 124,4393–4404 (1997).

14 Towers, M., Signolet, J., Sherman, A., Sang, H. & Tickle, C. Insights into bird wing evolution and digit specification from polarizing region fate maps. Nat Commun 2, 426 (2011).

15 Towers, M., Mahood, R., Yin, Y. & Tickle, C. Integration of growth and specification in chick wing digit-patterning. Nature 452,882–886 (2008).

16 Tickle, C. & Towers, M. Sonic Hedgehog Signaling in Limb Development. Frontiers in cell and developmental biology 5, 14, doi:10.3389/fcell.2017.00014 (2017).

17 Tickle, C., Summerbell, D. & Wolpert, L. Positional signalling and specification of digits in chick limb morphogenesis. Nature 254,199–202 (1975).

18 Pickering, J. & Towers, M. Inhibition of Shh signalling in the chick wing gives insights into digit patterning and evolution. Development 143,3514–3521 (2016).

19 McKinnell, I. W., Turmaine, M. & Patel, K. Sonic Hedgehog functions by localizing the region of proliferation in early developing feather buds. Dev Biol 272,76–88, doi:10.1016/j.ydbio.2004.04.019 (2004).

20 Harris, M. P., Williamson, S., Fallon, J. F., Meinhardt, H. & Prum, R. O. Molecular evidence for an activator-inhibitor mechanism in development of embryonic feather branching. Proc Natl Acad Sci U S A 102,11734–11739, doi:10.1073/pnas.0500781102 (2005).

21 Harris, M. P., Fallon, J. F. & Prum, R. O. Shh-Bmp2 signaling module and the evolutionary origin and diversification of feathers. J Exp Zool 294,160–176, doi:10.1002/jez.10157 (2002).

22 Grieshammer, U., Minowada, G., Pisenti, J. M., Abbott, U. K. & Martin, G. R. The chick limbless mutation causes abnormalities in limb bud dorsal-ventral patterning: implications for the mechanism of apical ridge formation. Development 122,3851–3861 (1996).

23 Wu, P. et al. Multiple Regulatory Modules Are Required for Scale-to-Feather Conversion. Mol Biol Evol 35,417–430, doi:10.1093/molbev/msx295 (2018).

24 Coumailleau, P. & Duprez, D. Sim1 and Sim2 expression during chick and mouse limb development. Int J Dev Biol 53,149–157, doi:10.1387/ijdb.082659pc (2009).

25 Michon, F., Forest, L., Collomb, E., Demongeot, J. & Dhouailly, D. BMP2 and BMP7 play antagonistic roles in feather induction. Development 135,2797–2805, doi:10.1242/dev.018341 (2008).

26 Prin, P. & Dhouailly, D. How and when the regional competence of chick epidermis is established: feathers vs. scutate and reticulate scales, a problem en route to a solution. Int. J. Dev. Biol 48,137–148 (2004).

27 Domyan, E. T. et al. Molecular shifts in limb identity underlie development of feathered feet in two domestic avian species. Elife 5, e12115, doi:10.7554/eLife.12115 (2016).

28 Wolpert, L. Positional Information and Pattern Formation. Curr Top Dev Biol 117,597–608, doi:10.1016/bs.ctdb.2015.11.008 (2016).

29 Aruga, J. et al. A novel zinc finger protein, zic, is involved in neurogenesis, especially in the cell lineage of cerebellar granule cells. J Neurochem 63,1880–1890 (1994).

30 Hamburger, V. & Hamilton, H. L. A series of normal stages in the development of the chick embryo. 1951. J. Morphol. 88,49–92 (1951).

